# A Chromosome End Without Terminal Telomere Repeats is Stable for Multiple Cell Divisions

**DOI:** 10.1101/2025.05.01.651670

**Authors:** Haitao Zhang, Julien Audry, Kurt W Runge

**Affiliations:** Department of Inflammation and Immunity, Lerner Research Institute, Cleveland Clinic Foundation, Cleveland, Ohio, 44195 USA; Department of Genetics and Genome Sciences, Case Western Reserve University, Cleveland, Ohio, 44106 USA; Nexelis, 525 Bd Cartier O, Laval, QC H7V 3S8, Canada

**Author notes:** Submitting and Corresponding author: Kurt W Runge. **Contributor Roles**. Conceptualization: KWR. Data curation: HZ. Formal analysis HZ and KWR. Funding acquisition: KWR. Investigation: HZ, JA, KWR. Methodology: HZ, JA. Project administration: KWR. Resources: JA, KWR. Validation: HZ. Visualization: HZ. Writing – original draft: KWR. Writing – review & editing: HZ and KWR.

## Abstract

We have formed new short telomeres in *Schizosaccharomyces pombe* using an inducible nuclease that cuts near telomere repeats in cells that lack, cannot recruit or cannot fully activate telomerase. Sequencing these new telomeres showed that cells can divide at least 4 times with ∼30 bp of non-telomeric sequence at the chromosome end in cells lacking telomerase, which contrasts with current models for the roles of terminal single-stranded telomere repeats and the telomere proteins in telomere protection and replication. Cells that cannot recruit or activate telomerase had similar results, with additional rearrangements or telomere repeat addition, respectively.

## Main text

Telomeres consist of tracts of short repeated sequences bound by a specialized protein complex called “shelterin” (de Lange 2018; Kim et al. 2017; Pan et al. 2015). This complex protects the chromosome end and recruits telomerase, which adds the short DNA repeats to the chromosome end to complete replication (Autexier and Lue 2006). Telomere DNA repeats consist of a double-stranded (ds) portion and a single-stranded (ss) 3’ extension. In *Schizosaccharomyces pombe*, the ds repeats are bound by Taz1 which recruits Rap1 and Poz1 (**Fig. 1A**)(Kanoh and Ishikawa 2001; Miller et al. 2005; Miyoshi et al. 2008). The ssDNA tail is bound by Pot1 in a sequence-specific manner, which protects the chromosome end (Baumann and Cech 2001; Baumann and Price 2010; Pitt and Cooper 2010), and recruits Tpz1 and Ccq1 that bridges the ss tail to the ds repeats (**Fig. 1A**). Ccq1 recruits telomerase through phosphorylation of Thr93, and Tpz1 has a role in activating telomerase as the Tpz1-K75A mutant allows telomerase association with the telomere, but addition of multiple new telomere repeats is blunted (Armstrong et al. 2014; Moser et al. 2011; Webb and Zakian 2012; Yamazaki et al. 2012). The proteins Stn1 and Ten1 bind to the 3’ ssDNA extension of telomere repeats to promote full replication of the telomere (Matmati et al. 2018; Puglisi et al. 2008; Takikawa et al. 2017).

**Figure 1.**
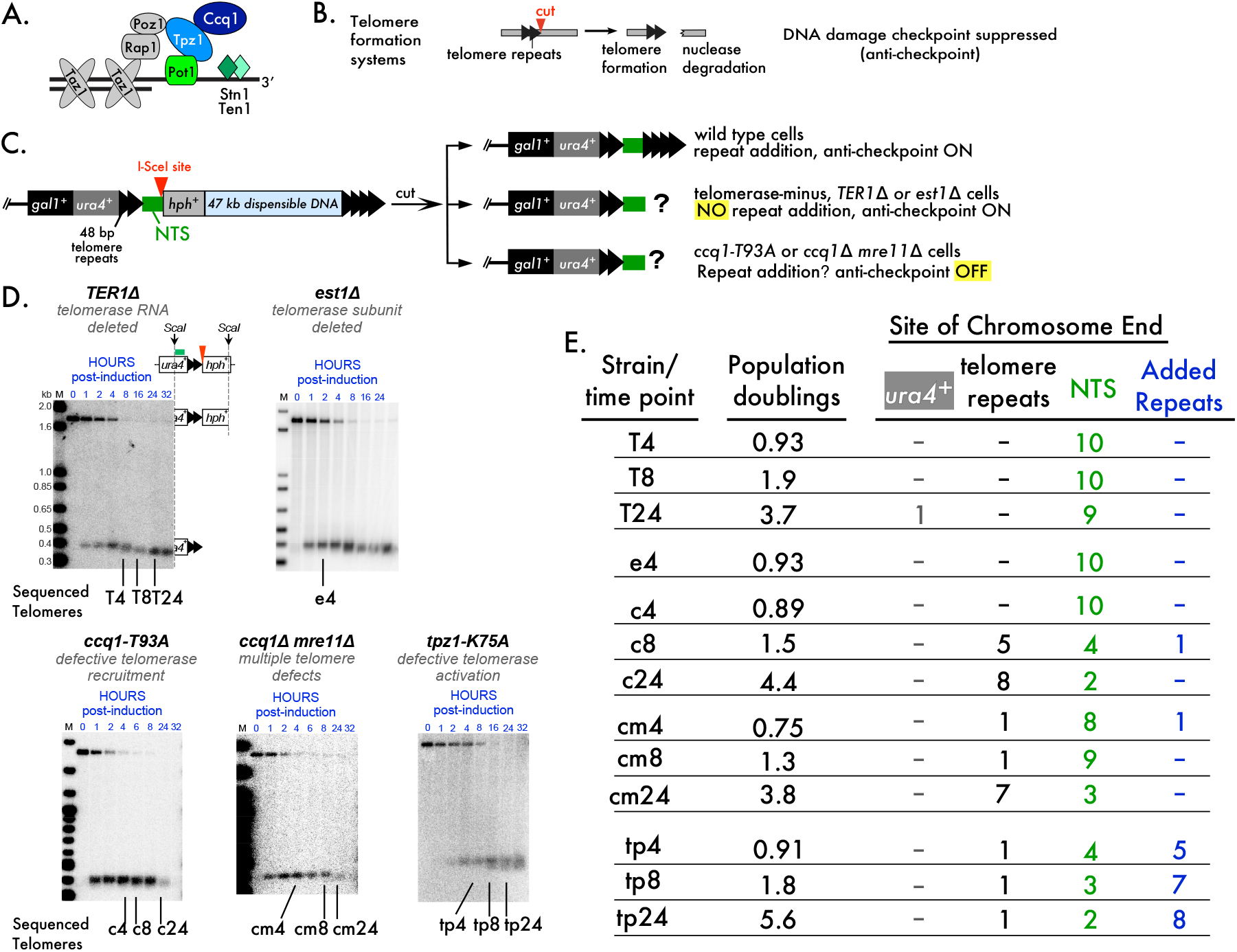
**A.** A schematic of *S. pombe* shelterin and the Stn1-Ten1 complex. The colored Ccq1- Tpz1-Pot1 complex is tethered to the single-stranded extension of telomere repeats. **B**. Telomere formation systems consist of a short tract of telomere repeats followed by several bp of sequence that include a restriction enzyme site that cuts only at the indicated site in the genome. The side bearing telomere repeats is elongated by telomerase while the other, non-telomeric, side is degraded. The telomere anti-checkpoint suppresses the DNA damage checkpoint that would be activated by degradation of the non-telomeric side. **C**. The telomere formation system used in this work (Audry et al. 2024; Wang et al. 2018). Production of I-SceI from an inducible promoter initiates cutting that is 90% complete in 4 h. Telomere repeat addition occurred at the non-telomeric sequence (NTS in green) in wild type cells with telomerase that have the anti-checkpoint, but repeat addition was not observed by Southern blotting in cells with telomere defects. **D**. Southern blots hybridized with a *ura4*^*+*^ probe to show the uncut and cut telomere formation system to monitor the fate of the new telomere (schematics of the uncut and cut telomere system are shown by the *TER1*Δ blot). The DNA timepoints used to clone and sequence telomeres are shown below each blot, where the first letter indicates the mutation and the second indicates the hour of the timepoint. The Southern blots were originally published in Audry et al., 2024 by Oxford University Press. **E**. Sites of telomere addition in the different mutants after different numbers of population doublings. The telomere formation system is divided into regions of *ura4*^*+*^, the 48 bp telomere repeats, the NTS and newly Added Repeats. Ten randomly cloned sequences for each timepoint, with no selection for size of insert, were sequenced. Alignments of these sequences are presented in the Extended Data.

Telomere formation systems in both *Saccharomyces cerevisiae* and *S. pombe* have used inducible nucleases that make unique cuts in chromatin adjacent to telomere repeats, leaving one end with telomere repeats that forms a new telomere and another that is degraded like a DNA double-strand break (DSB)(Bairley et al. 2011; Diede and Gottschling 1999; Wang et al. 2018)(**Fig. 1B**). The degraded DNA does not pause the cell cycle the way a DSB does (Audry et al. 2024; Michelson et al. 2005), indicating that the nearby telomere repeats cause an “anti- checkpoint” that suppresses the DNA damage response. In contrast, a DSB made in the same system is degraded in the first cell cycle and causes a cell cycle pause.

These telomere formation systems also revealed that telomere repeat addition occurs at non-telomeric DNA sequences adjacent to the nuclease cut (**Fig. 1C**, green block), and not in the telomere repeats themselves. These elongated telomeres are stable chromosome ends for multiple cell divisions (Bairley et al. 2011; Diede and Gottschling 1999; Wang et al. 2018).

We created an inducible system for forming telomeres and DSBs in *S. pombe* (Wang et al. 2018). The telomere formation system revealed that the shelterin protein Ccq1 is required to distinguish a short telomere from a DSB by preventing the nuclease Mre11 from degrading the new telomere (Audry et al. 2024). Ccq1 is phosphorylated at Thr93 to recruit telomerase to the telomere end for telomere repeat addition (Moser et al. 2011; Webb and Zakian 2012; Yamazaki et al. 2012). However, eliminating telomerase activity by eliminating the telomerase RNA (*TER1*Δ cells) or a protein subunit (*est1*Δ cells), eliminating telomerase recruitment with *ccq1* mutants (*ccq1-T93A* mutant or *ccq1*Δ *mre11*Δ mutant that delays DNA degradation) or inhibiting full telomerase activation at the telomere with the *tpz1-75A* mutation allowed the formation of stable chromosome ends. Thus, telomerase activity was not required to distinguish a new short telomere from a DSB (Audry et al. 2024; Wang et al. 2018).

A surprising observation was that the mutants lacking telomerase (*TER1*Δ, *est1*Δ) maintained a short, stable chromosome end for more than 24 h of growth or 3.7 population doublings (**Fig. 1D**). As I-SceI cuts >90% of its target sites within 4 h, these mutants have replicated the new end through several divisions (Audry et al. 2024). Similar results were obtained with other mutants defective in telomerase activity, *ccq1-T93A, ccq1*Δ *mre11*Δ and *tpz1-K75A* cells (**Fig. 1D**). Even though the *ccq1-T93A, ccq1*Δ *mre11*Δ and *tpz1-K75A* mutants express telomerase, the Southern blot results suggested that cells went through multiple divisions without noticeably elongating the I-SceI cut end.

These results raised questions regarding the nature of the new, stable telomere end. I-SceI cleavage produces an end with 31 bp of non-telomeric sequence (green NTS in **Fig. 1C**), the site of telomere addition in wild type cells (Wang et al. 2018). The new end could terminate in telomere repeats or the NTS (**Fig. 1C**). Preservation of the NTS would mean that the sequence-specific, single-strand DNA binding protein Pot1 would lack its normal binding site, as would the Stn1-Ten1 proteins (**Fig. 1A**)(Baumann and Price 2010; Puglisi et al. 2008; Takikawa et al. 2017). End stabilization via a t-loop structure, where the single-stranded telomere repeats invade and base pair with the double-strand repeats as seen in mammalian telomeres, would be highly unlikely with a NTS at the end (Griffith et al. 1999; Tomaska et al. 2020).

To resolve this question, we used the DNAs from different timepoints to sequence the newly formed stable ends using telomere PCR (**Fig. 1D**)(Audry et al. 2024). The telomere PCR approach adds oligo dC residues to DNA 3’ ends, allowing a specific primer in the telomere adjacent *ura4*^*+*^; DNA and an oligo dG primer to amplify the telomere end for cloning and sequencing (Förstemann et al. 2000). The oligo dC sequence identifies the 3’ end of the telomere DNA strand, and the sequence determines if the *ura4*^*+*^, 48 bp telomere repeats and NTS sequences are present and unrearranged. Ten telomere PCR clones, regardless of insert size, were isolated and sequenced for each of the timepoints tested (**Fig. 1D**). In this system, almost 100% of cells survive I-SceI cleavage and continue to grow, allowing 10 sequences to form a representative sample of the formed ends (Audry et al. 2024; Wang et al. 2018).

A striking result was that 39 out of 40 the new ends from cells lacking telomerase had some or all of the NTS at the chromosome end (**Fig. 1E, Extended data**). The single telomere that did not end with the NTS and had lost the 48 bp of telomere repeats was from cells that had grown for 3.7 population doublings after I-SceI cutting was induced, raising the possibility that this chromosome end was in the process of being degraded when the DNA was prepared.

The Ccq1 mutants that impair telomerase recruitment, lack the anti-checkpoint and retain telomerase activity gave different results. Some cells grown for ∼4 population doublings after cleavage was induced retained the NTS at the chromosome end, indicating that this unusual end was also stable in these mutants. Other telomeres had lost the NTS and had chromosomal termini that ended in the telomere repeats, and two of the telomeres had 10 or 12 bases of added telomere repeats (**Fig. 1E, Extended data**).

The *tpz1-K75A* mutant cells, which do not activate bound telomerase, grown for >5 doublings had new ends capped by the NTS and ends capped by the telomere repeats. Telomeres with three to 41 bases of added telomere repeats were isolated from these telomerase and anti- checkpoint positive cells (**Fig. 1E, Extended data**).

These results indicate that the “hybrid-end” consisting of a short stretch of telomere repeats capped by up to 33 bp of non-telomeric sequence can form a stable telomere in *S. pombe* for several cell divisions. This hybrid-end was most frequently isolated in cells that lack telomerase, and was still seen in cells with telomerase that have other defects in telomere addition. The lack of the telomere anti-checkpoint or inhibition of full telomerase activity did not prevent the stable maintenance of the hybrid-end over three to five population doublings.

The hybrid-end telomere formed after I-SceI cutting is immediately stable in the first cell cycle (Audry et al. 2024), indicating that cells must have a mechanism to transiently stabilize this structure as soon as the end forms. While chromosome ends capped by non-telomeric sequences have been identified in *S. cerevisiae* and *S. pombe* mutants (Jain et al. 2010; Maringele and Lydall 2005), these telomeres evolved after extensive genomic rearrangements and most likely represent a different pathway. Pot1 is an unlikely candidate to stabilize the newly formed hybrid end because Pot1 is not detected at the telomere repeats before I-SceI cleavage (Audry et al. 2024), and the hybrid end lacks Pot1 binding sites.

We hypothesize that the factor(s) that immediately stabilize the new hybrid-end for multiple population doublings normally play roles at telomeres with very short repeat tracts, stabilizing dysfunctional telomeres to allow telomerase elongation or cells to exit the cell cycle. Potential candidates for stabilizing the new hybrid-end include the DSB repair protein Rad52, which plays an important role at telomeres that become dysfunctional due to loss of telomere repeats (Abdallah et al. 2009; Khadaroo et al. 2009). The DNA end-binding complex Ku that also binds to telomeres and DSBs is another candidate (Baumann and Cech 2000; Gravel et al. 1998). The telomere formation system provides a means for testing a role for these proteins or identifying new players.

## Methods and Reagents

### Telomere PCR and sequencing

This method is modified from Förstermann et al. (Förstemann et al. 2000) using genomic DNA samples from (Audry et al. 2024). For the *est1*Δ experiment, only the 4 h timepoint had sufficient DNA for analysis. Genomic DNA from the lanes indicated in Fig. 1D was quantified with a Qubit 4 Fluorometer (ThermoFisher Scientific), and 500 ng was brought up to 34.5 μL with MilliQ-purified H_2_O, denatured at 96°C for 10 minutes and quickly cooled to 4°C for 20 minutes. The denatured sample was mixed with 5 μL of 2.5 mM CoCl_2_, 5 μL of 10X TdT buffer, 5 μL of 10 mM dCTP and 0.5 μL of Terminal Transferase (20 units/ μL, NEB catalog # M0315S) and incubated at 37°C for 30 minutes, followed by inactivation at 70°C for 10 minutes.

Tailing product (10 µL) was directly mixed with 25 μL of 2X DreamTaq Master Mix (ThermoFisher Scientific, catalog #: K1071), 1 μL of each primer u4-teloPCR-1S (10 μM, 5’- CCAATGAAAGATGTATGTAGATGAATG-3’) and BamHI-G18 (10 μM, 5’- CGGGATCCGGGGGGGGGGGGGGGGGG-3’), and 13 μL of MilliQ H_2_O. The PCR reaction was performed with an initial denaturation at 95 °C for 2 minutes, followed by 45 cycles of denaturation at 95 °C for 30 seconds, annealing at 56 °C for 30 seconds, and extension at 72 °C for 30 seconds, and a final extension at 72 °C for 10 minutes. The PCR product was then purified using the QIAquick PCR Purification Kit (catalog #28104).

The purified PCR product was TA-cloned into pCR2.1 vector (ThermoFisher Scientific, catalog #450641), transformed into TOP10 competent cells (ThermoFisher Scientific, catalog #C404006) and plated onto LB plates containing 50 μg/mL Kanamycin. The LB plates were pre-spread with 80 μL of 20 mg/mL X-gal (ThermoFisher Scientific, catalog #R0404) in dimethylformamide for blue-white screening. Fifteen colonies of each time point were randomly picked and cultured for plasmid preparation. The plasmids were randomly assigned numbers from 1 to 15. The first ten plasmids confirmed to contain inserts by EcoR I digestion were sequenced using the universal primers M13F and M13R.

The DNA sequences from each sequenced sample at each time point, spanning from the 5’ end of u4-teloPCR-1S binding site to the 3’ end where poly C tail was added, were aligned using the online software MAFFT version 7 (https://mafft.cbrc.jp/alignment/server/, accessed on April 16, 2025).

### Population doublings

The values in Fig. 1E were calculated by dividing the time after induction by the doubling times for each experiment determined in (Audry et al. 2024).

## Strains

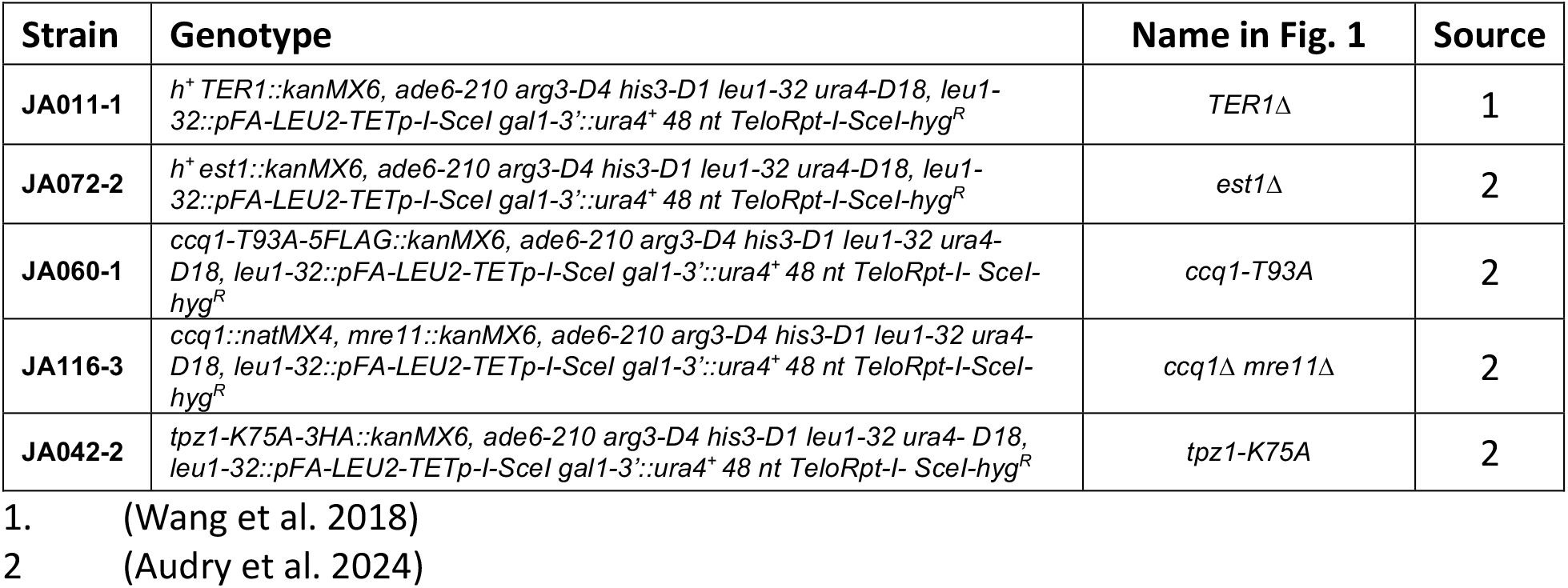

## Acknowledgements

We thank Dr. Bibo Li for helpful comments on the manuscript and Ms. Carly Kerr for technical assistance.

---

**Extended Data Figure 1.** The sequences for each timepoint are presented with clone numbers and sequences. Clone numbers are not consecutive as some clones did not have an insert. The “Ref.” sequence on the first line shows part of the 48 bp of telomere repeats (black font) and the 35 bp non-telomeric sequence that ends at the ATAA I-SceI 3’ overhang (green font). Newly added telomere repeats in bold italic blue font.

