## Extended Data for "A Chromosome End Without Terminal Telomere Repeats is Stable for Multiple Cell Divisions"

### Extend Data – sequence alignments

70 36 30 20 10 1
Ref. TACGGTTACACGGTTACACGGTTACAGGTTACAGGGGATCCGTCGACCTGCAGAGTTACGCTAGGGATAA

**TER1Δ 2R-48bp_JA011-1--4 hours**
Ref. TACGGTTACACGGTTACACGGTTACAGGTTACAGGGGATCCGTCGACCTGCAGAGTTACGCTAGGGATAA
1 TACGGTTACACGGTTACACGGTTACAGGTTACAGGGGATCCGTCGACCTGCAGAGTTACGCTAGGGATA
2 TACGGTTACACGGTTACACGGTTACAGGTTACAGGGGATCCGTCGACCTGCAGAGTTACGCTAGGGATAA
3 TACGGTTACACGGTTACACGGTTACAGGTTACAGGGGATCCGTCGACCTGCAGAGTTACGCTAGGGATAA
4 TACGGTTACACGGTTACACGGTTACAGGTTACAGGGGATCCGTCGACCTGCAGAGTTACGCTAGGG
5 TACGGTTACACGGTTACACGGTTACAGGTTACAGGGGATCCGTCGACCTGCAGAGTTACGCTAGGGATAA
7 TACGGTTACACGGTTACACGGTTACAGGTTACAGGGGATCCGTCGACCTGCAGAGTTACGCTAGGGATAA
8 TACGGTTACACGGTTACACGGTTACAGGTTACAGGGGATCCGTCGACCTGCAGAGTTACGCTAG
9 TACGGTTACACGGTTACACGGTTACAGGTTACAGGGGATCCGTCGACCTGCAGAGTTACGCTAGGGATAA
10 TACGGTTACACGGTTACACGGTTACAGGTTACAGGGGATCCGTCGACCTGCAGAGTTACGCTAGGGATAA
11 TACGGTTACACGGTTACACGGTTACAGGTTACAGGGGATCCGTCGACCTGCAGAGTTACGCTAGGGATAA

**TER1Δ 2R-48bp_JA011-1--8 hours**
Ref. TACGGTTACACGGTTACACGGTTACAGGTTACAGGGGATCCGTCGACCTGCAGAGTTACGCTAGGGATAA
1 TACGGTTACACGGTTACACGGTTACAGGTTACAGGGGATCCGTCGACCTGCAGAGTTACGCTAGGGATAA
2 TACGGTTACACGGTTACACGGTTACAGGTTACAGGGGATCCGTCGACCTGCAGAGTTACGCTAGGGATAA
3 TACGGTTACACGGTTACACGGTTACAGGTTACAGGGGATCCGTCGACCTGCAGAGTTACGCTAGGGATAA
4 TACGGTTACACGGTTACACGGTTACAGGTTACAGGGGATCCGTCGACCTGCAGAGTTACGCTAGGGATAA
5 TACGGTTACACGGTTACACGGTTACAGGTTACAGGGGATCCGTCGACCTGCAGAGTTACGCTAGGGATAA
6 TACGGTTACACGGTTACACGGTTACAGGTTACAGGGGATCCGTCGACCTGCAGAGTTACGCTAGGGATAA
7 TACGGTTACACGGTTACACGGTTACAGGTTACAGGGGATCCGTCGACCTGCAGAGTTACGCTAGGGATAA
8 TACGGTTACACGGTTACACGGTTACAGGTTACAGGGGATCCGTCGACCTGCAGAGTTACGCTAGGGATAA
9 TACGGTTACACGGTTACACGGTTACAGGTTACAGGGGATCCGTCGACCTGCAGAGTTACGCTAGGGATAA
10 TACGGTTACACGGTTACACGGTTACAGGTTACAGGGGATCCGTCGACCTGCAGAGTTACGCTAGGGATAA

**TER1Δ 2R-48bp_JA011-1--24 hours**
Ref. TACGGTTACACGGTTACACGGTTACAGGTTACAGGGGATCCGTCGACCTGCAGAGTTACGCTAGGGATAA
1 -122nt ----------------------------------------------------------------------------------------------------------------------------------------
2 TACGGTTACACGGTTACACGGTTACAGGTTACAGGGGATCCGTCGACCTGCAGAGTTACGCTAGGGATAA
3 TACGGTTACACGGTTACACGGTTACAGGTTACAGGGGATCCGTCGACCTGCAGAGTTACGCTAGGGA
4 TACGGTTACACGGTTACACGGTTACAGGTTACAGGGGATCCGTCGACCTGCAGAGTTACGCTAGGGATAA
5 TACGGTTACACGGTTACACGGTTACAGGTTACAGGGGATCCGTCGACCTGCAGAGTTACGCTAG
7 TACGGTTACACGGTTACACGGTTACAGGTTACAGGGGATCCGTCGACCTGCAGAGTTACGCTAGGGATAA
8 TACGGTTACACGGTTACACGGTTACAGGTTACAGGGGATCCGTCGACCTGCAGAGTTACGCTAG
9 TACGGTTACACGGTTACACGGTTACAGGTTACAGGGGATCCGTCGACCTGCAGAGTTACGCTA
10 TACGGTTACACGGTTACACGGTTACAGGTTACAGGGGAT
11 TACGGTTACACGGTTACACGGTTACAGGTTACAGGGGATCCGTCGACCTGCAGAGTTACGCTAGGGATAA

**est1Δ 2R-48 bp_JA072-2-4 hours**
Ref. TACGGTTACACGGTTACACGGTTACAGGTTACAGGGGATCCGTCGACCTGCAGAGTTACGCTAGGGATAA
1 TACGGTTACACGGTTACACGGTTACAGGTTACAGGGGATCCGTCGACCTGCAGAGTTACGCTAGGGATA

3 TACGGTTACACGGTTACACGGTTACAGGTTACAGGGGATCCGTCGACCTGCAGAGTTACGCTAGGGATAA
4 TACGGTTACACGGTTACACGGTTACAGGTTACAGGGGATCCGTCGACCTGCAGAGTTACGCTAGGGATAA
5 TACGGTTACACGGTTACACGGTTACAGGTTACAGGGGATCCGTCGACCTGCAGAGTTACGCTAGGGAT
6 TACGGTTACACGGTTACACGGTTACAGGTTACAGGGGATCCGTCGACCTGCAGAGTTACGCTAGGGATAA
7 TACGGTTACACGGTTACACGGTTACAGGTTACAGGGGATCCGTCGACCTGCAGAGTTACGCTAGGGATA
10 TACGGTTACACGGTTACACGGTTACAGGTTACAGGGGATCCGTCGACCTGCAGAGTTACGCTAGGGATAA
11 TACGGTTACACGGTTACACGGTTACAGGTTACAGGGGATCCGTCGACCTGCAGAGTTACGCTAGGGACA
12 TACGGTTACACGGTTACACGGTTACAGGTTACAGGGGATCCGTCGACCTGCAGAGTTACGCTAGGG
13 TACGGTTACACGGTTACACGGTTACAGGTTACAGGGGATCCGTCGACCTGCAGAGTTACGCTAGGGATAA

**ccq1-T93A-FLAG 2R-48 bp_JA60-1--4 hours**

Ref. TACGGTTACACGGTTACACGGTTACAGGTTACAGGGGATCCGTCGACCTGCAGAGTTACGCTAGGGATAA
1 TACGGTTACACGGTTACACGGTTACAGGTTACAGGGGATCCGTCGACCTGCAGAGTTACGCTAGGGATAA
2 TACGGTTACACGGTTACACGGTTACAGGTTACAGGGGATCCGTCGACCTGCAGAGTTACGCTAGGGACAA
3 TACGGTTACACGGTTACACGGTTACAGGTTACAGGGGATCCGTCGACCTGCAGAGTTACGCTAG
4 TACGGTTACACGGTTACACGGTTACAGGTTACAGGGGACCCGTCGACCTGCAGAGTTACGCTAGGGATAA
5 TACGGTTACACGGTTACACGGTTACAGGTTACAGGGGATCCGTCGACCTGCAGAGTTACGCTAGGGA
6 TACGGTTACACGGTTACACGGTTACAGGTTACAGGGGATCCGTCGACCTGCAGAGTTACGCTAGG
7 TACGGTTACACGGTTACACGGTTACAGGTTACAGGGGATCCGTCGACCTGCAGAGTTACG
8 TACGGTTACACGGTTACACGGTTACAGGTTACAGGGGATCCGTCGACCTGCAGAGTTACGCTAGGGAT
9 TACGGTTACACGGTTACACGGTTACAGGTTACAGGGGATCCGTCGACCTGCAGAGTTACGCTAGGGATAA
10 TACGGTTACACGGTTACACGGTTACAGGTTACAGGGGATCCGTCGACCTGCACAGTTACCCTAGGGATAA

**ccq1-T93A-FLAG 2R-48 bp_JA60-1--8 hours**

Ref. TACGGTTACACGGTTACACGGTTACAGGTTACAGGGGATCCGTCGACCTGCAGAGTTACGCTAGGGATAA
1 TACGGTTACACGGTTACACGGTTACAGGTTACAGGGGATCCGTCGACCTGCAGAGTTACGCTAGGGATA
2 TACGGTTACACGGTTACACGGTTACAGGTTACAGGGGATCCGTCGACCTGCAGAGTTACGCTAGGGATTA***CGGTTACAGG***
3 TACGGTTACACGGTTACACGGTTACAGGTTACAGGGGATCCGTCGACCTGCAGAGTTACG
4 TACGGTTACACGGTTACA
5 TACGGTTACACGGTTACACGGTTACAGGTTACAGGGGATCCGTCGACCTGCAGAGTTA
6 TACGGTTACACGGTTACACGGTTACAGGTTACAG
7 TACGGTTACACGGTTACACGGTTACAGGTTACAGGGGATCCGTCGACCTGCAGAGTTACGCTAGGGATAA
8 TACGGTTACACGGTTACA
9 TACGGTTACACGGTTACACGGTTACAG
10 TACGGTTACACGGTTACA

**ccq1-T93A-FLAG 2R-48 bp_JA60-1--24 hours**

Ref. TACGGTTACACGGTTACACGGTTACAGGTTACAGGGGATCCGTCGACCTGCAGAGTTACGCTAGGGATAA
1 TACGGTTACACGGTTACACGGTTACAGGTTACA
2 TACGGTTACACGGTTACGCGGTTA
4 TACGGTTACACGGTTACACGGTTACAGGTTACAGGGGATCCGTTGACCTGCAGAGGTTACGGTTA
5 TACGGTTACACGGTTACACGGTTACAGGTTACAGGGGAT
6 TACGGTTACACGGTTACACGGTTACAGGTTACAG
7 TACGGTTACACGGTTACACGGTTACA
8 TACGGTTACACGGTTACACGGTTACAGGTTACAG
10 TACGGTTACACGGTTACACA
11 TACGGTTACACGGTTACACGGTTACAGGTTA
12 TACGGTTACACGGTTACACGGTTACAGGTTACAG

**ccq1Δ mre11Δ 2R-48 bp_JA116-3--4hours**

Ref. TACGGTTACACGGTTACACGGTTACAGGTTACAGGGGATCCGTCGACCTGCAGAGTTACGCTAGGGATAA
1 TACGGTTACACGGTTACACGGTTACAGGTTACAGGGGATCCGTCGACCTGCAGAGTTACGCTAGGGATAA***CGGTTACGGTTA***
2 TACGGTTACACGGTTACACGGTTACAGGTTACAGGGGATCCGCCGACCTGCAGAGTTACGCTAGGGATAA
3 TACGGTTACACGGTTACACGGTTACAGGTTACAGGGGATCCGTCGACCTGCAGAGTTACGCTAGGGATAA
4 -79nt-----------------------------------------------------------------------------------------------------------------------------
6 TACGGTTACACGGTTACACGGTTACAGGTTACAGGGGATCCGTCGACCTGCAGAGTTACGCTAGGGA
7 TACGGTTACACGGTTACACGGTTACAGGTTACAGGGGATCCGTCGACCTGCAGAGTTACGCTAGGGATA
8 TACGGTTACACGGTTACACGGTTACAGGTTACAGGGGATCCGTCGACCTGCAGAGTTACGCTAGGGATAA
9 TACGGTTACACGGTTACACGGTTACAGGTTACAGGGGATCCATCGACCTGCAGAGTTACGCTAGG
10 TACGGTTACACGGTTACACGGTTACAGGTTACAGGGGATCCGTCGACCTGCAGAGTTACGCTAGGGATAA
11 TACGGTTACACGGTTACACGGTTACAGGTTACAGGGGATCCGTCGACCTGCAGAGTTACGCTAGGGATAA

**ccq1Δ mre11Δ 2R-48 bp_JA116-3--8hours**

Ref. TACGGTTACACGGTTACACGGTTACAGGTTACAGGGGATCCGTCGACCTGCAGAGTTACGCTAGGGATAA
1 TACGGTTACACGGTTACACGGTTACAGGTTACAGGGGATCCGTCGACCTGCAGAGTTACGCTAGGGA
2 TACGGTTACACGGTTA
4 TACGGTTACACGGTTACACGGTTACAGGTTACAGGGGATCCGTCGACCTGCAGAGTTACGCTAG
5 TACGGTTACACGGTTACACGGTTACAGGTTACAGGGGATCCGTCGACCTGCAGAGTTACGCTAGGG
6 TACGGTTACACGGTTACACGGTTACAGGTTACAGGGGATCCGTCGACCTGCAGAGTTACGCTAGGGATAA
7 TACGGTTACACGGTTACACGGTTACAGGTTACAGGGGATCCGTCGACCTGCAGAGTTACGCTAGGGATAA
8 TACGGTTACACGGTTACACGGTTACAGGTTACAGGGGATCCGTCGACCTGCAGAGTTACGCTAGGGATAA
9 TACGGTTACACGGTTACACGGTTACAGGTTACAGGGGATCCATCGACCTGCAGAGTTACGCTAGGG
11 TACGGTTACACGGTTACACGGTTACAGGTTACAGGGGATCCGTCGACCTGCAGAGTTACGCTAGGGATA
12 TACGGTTACACGGTTACACGGTTACAGGTTACAGGGGATCCGTCGACCTGCAGAGTTACGCTAGG

**ccq1Δ mre11Δ 2R-48 bp_JA116-3--24hours**

Ref. TACGGTTACACGGTTACACGGTTACAGGTTACAGGGGATCCGTCGACCTGCAGAGTTACGCTAGGGATAA
1 TACGGTTACACGGTTACACGGTTACAG
2 TACGGTTACACGGTTACA
3 TACGGTTACACGGTTACACGGTTACAGGTTACAG
4 TACGGTTACACGGTTACACGGTTACAGGTTACAGGG
5 TACGGTTACACGGTTACACGGTTACAGGTTACAG
6 TACGGTTACACGGTTACACGGTTACAGGTTACAG
7 TACGGTTACACGGTTACACGGTTACAGGTTACAGGGGATCCGTCGACCTG
8 TACGGTTACACGGTTACACGGTTACAGGTTACAG
9 TACGGTTACACGGTTACACGGTTACAGGTTACAGGGGATCCGTCGACCTG
10 TACGGTTACACGGTTACACGGTTACAGGTTACAGG

**Tpz1-K75A-HA 2R-48 bp_JA042-2--4 hours**

Ref. TACGGTTACACGGTTACACGGTTACAGGTTACAGGGGATCCGTCGACCTGCAGAGTTACGCTAGGGATAA
1 TACGGTTACACGGTTACACGGTTACAGGTTACAGGGGATCCGTCGACCTGCAGAGTTACGCTAGGGATAA
2 TACGGTTACACGGTTACACGGTTACAGGTTACAGGGGATCCGTCGACCTGCAGAGTTACG
3 TACGGTTACACGGTTACACGGTTACAGGTTACAGGGGACCCGTCGACCTGCAGAGTTACGCTAGGGA
4 TACGGTTACACGGTTACACGGTTACAGGTTACAGGGGATCCGTCGACCTGCAGAGTTACGCTAGG***GGTTA***
5 TACGGTTACACGGTTA
6 TACGGTTACACGGTTACACGGTTACAGGTTACAGGGGATCCGTCGACCTGCAGAGTTACGCTAGGGATAA***CGGT***
7 TACGGTTACACGGTTACACGGTTACAGGTTACAGGGGATCCGTCGA
9 TACGGTTACACGGTTACACGGTTACAGGTTACAGGGGATCCGTCGACCTGCAGAGTTACGC***TTACACGGTTACAGGGTTACA***

10 TACGGTTACACGGTTACACGGTTACAGGTTACAGGGGATCCGTCGACCTGCAGAGTTACGCTAGGGATA***CGGTTACAGGTTACAG***
11 TACGGTTACACGGTTACACGGTTACAGGTTACAGGGGATCCGTCGACCTGCAGAGTTACGCTAGGG***GTTACAG***

**Tpz1-K75A-HA 2R-48 bp_JA042-2--8 hours**

Ref. TACGGTTACACGGTTACACGGTTACAGGTTACAGGGGATCCGTCGACCTGCAGAGTTACGCTAGGGATAA
1 TACGGTTACACGGTTACACGGTTACAGGTTA
2 TACGGTTACACGGTTACACGGTTACAGGTTACAGGGGATCCGTCCACCTGCCCAGTTACGCTAGG***TTACAGGGGTTACGGTTA***
3 TACGGTTACACGGTTACACGGTTACAGGTTACAGGGG***TTA***
4 TACGGTTACACGGTTACACGGTTACAGGTTACAGGGGATCCGTCGACCTGCAGAGTTACGCTAGGGATAA***CGGTTA***
5 TACGGTTACACGGTTACACGGTTACAGGTTACAGGGGAT
6 TACGGTTACACGGTTACACGGTTACAGGTTACAGGGGATCCGTCGACCTGCAGAGTTACGCTAGGGG***TTACGGTTACA***
7 TACGGTTACACGGTTACACGGTTACAGGTTACAGGGGATCCGTCGACCTGCAGAGTTACGCTAGGGATA***CGG***(***TTACAGGG)***$_{30nt}$
8 TACGGTTACACGGTTACACGGTTACAGGTTACAGGGGA
9 TACGGTTACACGGTTACACGGTTACAGGTTACAGGGGATCCGTCGACCTGCAGGGTTACGCTAGGGATAA***GGTTACAGGGG***
10 TACGGTTACACGGTTACACGGTTACAGGTTACAGGGGATCCGTCGACCTGCAGAGTTACGCTAGGGATA***CGGTTACGGTTACAGGTTA***

**Tpz1-K75A-HA 2R-48 bp_JA042-2--24 hours**

Ref. TACGGTTACACGGTTACACGGTTACAGGTTACAGGGGATCCGTCGACCTGCAGAGTTACGCTAGGGATAA
1 TACGGTTACACGGTTACACGGTTACAGGTTACAGGGGATCCGTCGACCTGCAGAGTTACGCTAGGGATA***CGGTTACAGG***
2 TACGGTTACACG
3 TACGGTTACACGGTTACACGGTTACAGGTTACAGGGGATCCGTCGACCTGCAGAGTTACGCTAGGGATA***CGG(TTACACGG)***$_{38nt}$
4 TACGGTTACACGGTTACACGGTTACAGGTTACAGG***TTA***
5 TACGGTTACACGGTTACACGGTTACAGGTTACAGGGGATCCGTCGACCTGCAGAGTTACGC***GGTTACAGGGTTTA***
6 TACGGTTACACGGTTACACGGTTACAGGTTACAGGGGATCCGTCGACCTGCAGAGTTACGCTAGG***TTACAGGTTACACGGTTA***
7 TACGGTTACACGGTTACACGGTTACAGGTTACAGGGGATCCGTCGACCTGCAGAGTTACGCTAGGGATAA***GGGTTACAGGGTACGGTTA***
8 TACGGTTACACGGTTACACGGTTACAGGTTACAGGGGATCCGTCGACCTGCAGAGTTACGCTAGGGATA***CGG(TTACAGGG)***$_{22nt}$
10 TACGGTTACACGGTTACACGGTTACAGGTTACAGGG
11 TACGGTTACACGGTTACACGGTTACAGGTTACAGGGGATCCGTCGACCTGCAGAGTTACGCTAGGG***TTACGG***
